# Molecular determinants of phase separation for *Drosophila* DNA replication licensing factors

**DOI:** 10.1101/2021.04.12.439485

**Authors:** Matthew W. Parker, Jonchee Kao, Alvin Huang, James M. Berger, Michael R. Botchan

## Abstract

Liquid-liquid phase separation (LLPS) of intrinsically disordered regions (IDRs) in proteins can drive the formation of membraneless compartments in cells. Phase-separated structures enrich for specific partner proteins and exclude others. We have shown that the IDRs of metazoan DNA replication initiators drive DNA-dependent phase separation *in vitro* and chromosome binding *in vivo*, and that initiator condensates selectively recruit specific partner proteins. How initiator IDRs facilitate LLPS and maintain compositional specificity is unknown. Using *D. melanogaster (Dm)* Cdt1 as a model initiation factor, we show that phase separation results from a synergy between electrostatic DNA-bridging interactions and hydrophobic inter-IDR contacts. Both sets of interactions depend on sequence composition (but not sequence order), are resistant to 1,6- hexanediol, and do not depend on aromaticity. These findings demonstrate that distinct sets of interactions drive self-assembly and condensate specificity across different phase-separating systems and advance efforts to predict IDR LLPS propensity and specificity *a priori*.

## INTRODUCTION

The internal environment of a cell is compartmentalized by both membrane-bound organelles and by protein-rich and protein/nucleic acid-rich compartments that lack an enclosing membrane. These membraneless compartments are commonly called biomolecular condensates and form through an ability of their constituent factors to undergo liquid-liquid phase separation (**LLPS**) (reviewed in (Banani et al., 2017)). Cellular bodies that form by LLPS are often spherical due to surface tension minimization across the phase boundary (Elbaum- Garfinkle, 2019). Sphericity, however, is not a defining feature of biomolecular condensates and many variables contribute to droplet (de)formation, including the physical parameters of the liquid phase (e.g., surface tension, viscosity and size) and external variables (e.g., application of a force and surface interactions). Non-spherical condensates often nucleate from or assemble along relatively rigid intracellular scaffolds. This effect is seen with cellular signaling complexes (Banjade and Rosen, 2014; Li et al., 2012; Su et al., 2016) and tight junctions (Beutel et al., 2019), which spread with the dimensions of the plasma membrane. Likewise, the chromatin-associated synaptonemal complex (Rog et al., 2017), the perichromosomal layer (Booth and Earnshaw, 2017) and metazoan DNA replication initiation proteins (Parker et al., 2019) are predicted to form a protein rich phase that contours with the more rigid nature of their substrate, mitotic chromosomes (Batty and Gerlich, 2019; Goloborodko et al., 2016; Houlard et al., 2015; Sun et al., 2018).

We previously discovered that the factors responsible for initiating DNA replication in metazoans, in particular the fly homologs of the Origin Recognition Complex (ORC) and Cdc6, as well as fly and human Cdt1, undergo DNA-dependent LLPS at physiological salt and protein concentrations *in vitro* (Parker et al., 2019). Recent studies demonstrate that human Orc1 can also phase separate in the presence of DNA (Hossain et al., 2021). In live cells, initiators first associate with chromatin in mitosis where they appear to uniformly coat, or “wet”, anaphase chromosomes (Baldinger and Gossen, 2009; Kara et al., 2015; Parker et al., 2019; Sonneville et al., 2012). These cytological observations are consistent with expectations for a protein that undergoes a condensation reaction with a relatively rigid intracellular scaffolding partner. The ability to condense on DNA is conferred by a metazoan-specific, N-terminal intrinsically disordered region (**IDR**) present in Orc1, Cdc6 and Cdt1. For Orc1 at least, this region is required for binding chromatin in tissue culture cells and for viability in flies (Parker et al., 2019). *In vitro* assays show that the Orc1 IDR is dispensable for ATP-dependent DNA-binding by *D. melanogaster* ORC and loading of the Mcm2-7 replicative helicase (Schmidt and Bleichert, 2020). However, at physiological ionic strength and protein concentrations initiator IDRs drive initiator coalescence into a condensed phase that supports ATP-dependent recruitment of Mcm2-7 and which can be controlled by CDK-dependent phosphorylation, an event that directly regulates initiator mechanism (Parker et al., 2019). Together, the available data suggest that LLPS facilitates the formation of an enriched layer of initiation factors that coats mitotic chromosomes to enhance the kinetics of helicase loading and enable complete replication licensing within a relatively short period of time (Dimitrova et al., 2002; Méndez and Stillman, 2000; Okuno et al., 2001).

How metazoan initiator IDRs facilitate DNA-dependent LLPS at a biophysical level is unknown. Protein multivalency underlies the formation of intermolecular interaction networks that can drive LLPS and, in many phase-separating systems, functional multivalency is contributed by the presence of one or more IDRs (Li et al., 2012). Although phase-separating IDR sequences vary between systems, some common sequence features capable of supporting LLPS have begun to emerge. For example, many phase-separating IDRs have low amino acid sequence complexity (so-called low complexity domains (**LCDs**)) and are highly enriched for a few select amino acids (e.g., the nucleolar protein FIB1 (Feric et al., 2016), the P-granule protein Laf-1 (Elbaum-Garfinkle et al., 2015), the stress granule protein hnRNPA1 (Molliex et al., 2015), the transcription factor EWS (Chong et al., 2018), and the transcriptional coactivator MED1 (Sabari et al., 2018)). The prevalence of low-complexity disordered sequences in phase-separating proteins has led to a commonly-held view that LCDs are a universal feature of IDR-mediated LLPS. However, there exist important exceptions to this correlation, including phase separation by the intrinsically disordered Nephrin Intracellular Domain (NICD) (Pak et al., 2016) and the metazoan replication initiators (Parker et al., 2019), both of which possess high sequence-complexity IDRs. Thus, LCDs are but a subtype of phase-separating IDR.

Literature reports of phase separating sequences are increasing rapidly and yet relatively few of these studies contain a biophysical description of LLPS. This gap leaves open the question of whether different biomolecular condensates form by a generally shared or individually distinctive set of cohesive interactions. Interestingly, many condensates (**Table 1**) dissolve upon treatment with the aliphatic alcohol 1,6-hexanediol (1,6-HD), a compound that permeabilizes the nuclear pore by disrupting weak hydrophobic Phe-Gly (‘FG’)-repeat interactions that are prevalent in the assembly (Patel et al., 2007; Shulga and Goldfarb, 2003). The broad sensitivity of protein LLPS to 1,6-HD could be taken to indicate that phase-separation mechanisms are generalizable, at least for certain IDR classes. Consistently, sequence aromaticity has been shown to be an essential feature in a variety of phase-separating systems (Chiu et al., 2020; Chong et al., 2018; Lin et al., 2017; Nott et al., 2015; Pak et al., 2016; Qamar et al., 2018; Wang et al., 2018). Despite these similarities, however, membraneless organelles *in vivo* have a particular compositional bias that often underpins their utility, implying that there exists a molecular ‘grammar’ or ‘sorting code’ that accepts a particular subset of partner factors while excluding inappropriate factors. Consistent with this idea, we have observed that initiator IDRs can co-recruit pathway-specific partner proteins but not other types of phase-separating proteins (such as FUS) (Parker et al., 2019). Our understanding of the molecular rules that govern IDR sorting mechanisms is still in its infancy.

**Table 1:** List of proteins with condensate sensitivity to 1,6-hexanediol.

| Protein | Reference(s) |
| --- | --- |
| TDP43 | (Babinchak et al., 2019; Schmidt et al., 2019) |
| hnRNPA1 | (Molliex et al., 2015) |
| Tau | (Wegmann et al., 2018) |
| Huntingtin protein exon 1 | (Peskett et al., 2018) |
| Rnq1 | (Kroschwald et al., 2015) |
| BRD4 | (Sabari et al., 2018) |
| MED1 | (Cho et al., 2018; Sabari et al., 2018) |
| Hp1 | (Strom et al., 2017) |
| hnRNPA2 | (Lin et al., 2016) |
| TACC3 | (So et al., 2019) |
| FUS | (Kroschwald et al., 2017) |
| RPB1 | (Boehning et al., 2018) |
| Chromosome passenger complex | (Trivedi et al., 2019) |
| Synaptonemal complex | (Rog et al., 2017) |

Here we use *Drosophila melanogaster* Cdt1 as a model to dissect the molecular basis for DNA- dependent LLPS by metazoan replication initiator proteins. We find that phase separation by Cdt1 is unaffected by treatment with 1,6-hexanediol and that sequence aromatics, which represent < 3% of the total initiator IDR amino acids, are dispensable for condensate formation. We show that protein-DNA interactions are electrostatic in nature and contribute an adhesive force to LLPS, with DNA functioning as a counterion bridge. These interactions synergize with cohesive intermolecular IDR-IDR interactions, which are driven by the hydrophobic effect. Using mutagenesis, we demonstrate that intermolecular IDR-IDR interactions, not DNA-mediated bridging interactions, serve as the preeminent driving force behind initiator phase separation. Collectively, these studies provide a detailed picture of the mechanism of initiator-type IDR LLPS and demonstrate that biomolecular condensates can form by a variety of different molecular interaction types.

## RESULTS & DISCUSSION

### Initiator liquid-liquid phase separation is resistant to treatment with 1,6-hexanediol and does not require aromatic residues

The IDRs of the metazoan replication initiation factors are necessary and sufficient for DNA-dependent LLPS (Parker et al., 2019). To understand the mechanism of initiator condensation, we first set out to identify similarities in amino acid sequence composition between initiator IDRs and the IDRs of other condensate forming proteins. We generated a sequence heatmap to compare the fractional representation of each amino acid in the IDRs of Orc1 (residues 187-549), Cdc6 (residues 1-246), and Cdt1 (residues 1-297) with the IDRs of a small suite of known condensate-forming proteins (human Fused in Sarcoma (FUS) (residues 1-293), *C. elegans* Laf- 1 (residues 1-203), human Ddx4 (residues 1-260), human hnRNPA1 (residues 178-372), human DNA-directed RNA polymerase II subunit RPB1 (residues 1531-1970), human Mediator of RNA polymerase II transcription subunit 1 (MED1) (residues 948-1574), and mouse BuGZ (residues 63-495)) (**Figure 1A**). This analysis revealed major differences in initiator IDR amino acid composition compared to the other proteins analyzed and demonstrate that initiator IDRs represent a sequence class all their own, which we term “initiator-type” IDRs. The most striking observation was the conspicuous absence of any one or more highly enriched residue within the initiator IDRs (dark red = fraction composition > 15%). All sequences in the comparison suite possess at least one highly enriched residue, most often serine, proline, and/or glycine. By contrast, initiator IDR composition is distributed more uniformly across all amino acids, with the exception of histidine, cysteine and tryptophan (which are weakly represented amongst all IDRs assessed), and methionine, glycine and aromatic residues (which are weakly represented in the initiators but not necessarily in the other IDRs). The broad utilization of amino acid sequence space by initiator IDRs suggested that these sequences have higher complexity than the other sequences analyzed. To quantitatively assess complexity, we calculated the informational entropy of each IDR sequence. Protein informational entropy, or sequence complexity, is the amount of information stored in a given linear sequence of amino acids and is calculated from the number of observed occurrences of each of the twenty amino acids (Wootton and Federhen, 1993). This analysis revealed that initiator IDRs have a higher sequence complexity than all other condensate-forming IDRs analyzed, except for Ddx4, which has a similar complexity score. The unique sequence signature of initiator IDRs was a first indication that initiator LLPS might proceed through a mechanism distinct from other phase-separating proteins.

**Figure 1:**
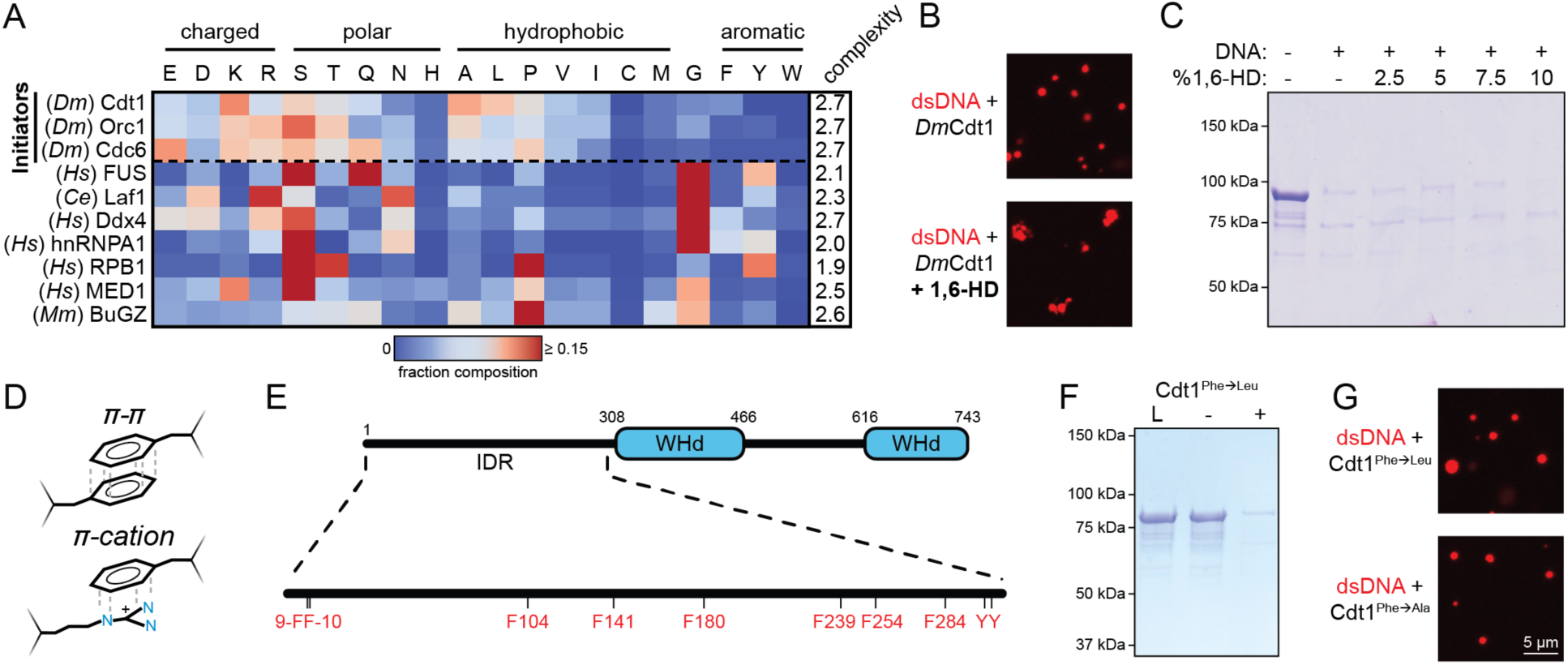
Initiator phase separation is insensitive to 1,6-hexanediol and does not require aromatic residues. A) Heatmap representation of the amino acid composition of the three *Drosophila* initiation factors (Cdt1, Orc1 and Cdc6) compared to other phase separating disordered sequences. B) Confocal fluorescence microscopy images of condensates formed with 5 µM Cy5-dsDNA and 5 µM Cdt1 in the absence (top) and presence (bottom) of 1,6- hexanediol. C) 2 µM Cdt1 was combined with 2 µM dsDNA and phase separation was assessed with a depletion assay in the presence of increasing concentrations (w/v) of 1,6-HD. Phase separation is indicated by protein loss. D) Aromatic residues contribute to the phase separation of many condensate systems through their ability to participate in π-π and π-cation interactions. E) Schematic of the *D. melanogaster* Cdt1 protein (top). Cdt1 contains two winged-helix domains (WHd, blue) and an N-terminal disordered domain (black line). The N-terminal IDR contains multiple aromatic residues (bottom). F) Depletion assay results assessing phase separation of the Cdt1 aromatic mutant, Cdt1^Phe→Leu^. For this construct all phenylalanine residues have been mutated to leucine. “L” is a load control, “-“ is in the absence of DNA, and “+” is in the presence of DNA. G) Confocal fluorescence microscopy images of condensates formed with 5 µM Cy5- dsDNA and either Cdt1^Phe→Leu^ (top) or Cdt1^PheàAla^ (bottom).

The aliphatic alcohol 1,6-hexanediol (1,6-HD) can inhibit the self-assembly and phase separation of compositionally distinct classes of disordered sequences (see **Table 1** references). This behavior suggests that even for different IDR sequences, similar molecular interactions are responsible for driving LLPS. We therefore tested whether condensates formed from initiator IDRs are likewise sensitive to 1,6-HD. Initiator LLPS was directly visualized by fluorescence microscopy by mixing ORC, Cdc6, and Cdt1 with a Cy5-labeled sixty base pair double-stranded DNA (Cy5-dsDNA). LLPS by all three proteins proved to be resistant to treatment with 10% 1,6- HD (**Figure 1B** and **Figure 1-figure supplement 1A-B**). Interestingly, 1,6-HD did lead to morphological changes in initiator/DNA condensates, including droplet clustering and, for ORC, a reduction in overall size. To confirm the microscopy results, we assessed the effect of 1,6-HD on Cdt1 partitioning into a condensed phase by a depletion assay. In this assay, LLPS is induced by the addition of sixty base pair dsDNA, the denser phase separated material is pelleted by centrifugation, and LLPS is assessed by protein depletion from the supernatant. Titrations of 1,6-HD from 0-10% (w/v) again showed no effect on Cdt1 LLPS (**Figure 1C**), confirming that initiator phase separation is insensitive to this reagent.

Aromatic residues play a critical role in driving protein LLPS, where they contribute multivalent π-π and π-cation interactions (**Figure 1D**) (Chiu et al., 2020; Chong et al., 2018; Lin et al., 2017; Nott et al., 2015; Pak et al., 2016; Qamar et al., 2018; Vernon et al., 2018; Wang et al., 2018). However, the resistance of initiator LLPS to treatment with 1,6-HD (**Figure 1B-C**), as well as the low (< 3%) aromatic residue content of initiator IDRs, suggested that these interactions may not be a primary driving force for phase separation. The IDR of Cdt1 contains only eight aromatic residues, all phenylalanine, that are relatively equally distributed throughout the region (**Figure 1E**). To test whether initiator LLPS is governed by aromatic-mediated interactions, we mutated all phenylalanines in the Cdt1 IDR to leucine (Cdt1^Phe→Leu^) and assessed phase separation by both the depletion assay (**Figure 1F**) and fluorescence microscopy (**Figure 1G**, top panel). The replacement of Phe with Leu abolishes aromaticity while maintaining a comparable level of hydrophobicity (phenylalanine hydrophobicity index = 2.8, leucine hydrophobicity index = 3.8, (Kyte and Doolittle, 1982)). Interestingly, the loss of aromaticity had no detectable effect on DNA-induced phase separation propensity. Similarly, mutating Cdt1’s aromatic residues to alanine (Cdt1^Phe→Ala^) also did not block phase separation (**Figure 1G**, bottom panel). Together, these data show that initiator phase separation does not rely on aromatic residue-mediated interactions, consistent with its ability to resist treatment with 1,6-HD.

### Initiator IDR electrostatics underly coacervation with polyanionic scaffolds

The DNA dependency of initiator LLPS suggested to us that initiator droplets form by coacervation, a process in which two oppositely charged polymers drive the formation of a condensed phase (Sing and Perry, 2020). We therefore set out to test the sensitivity of Cdt1 LLPS to salt. Using the depletion assay, we measured phase separation at different concentrations of potassium glutamate (KGlu; 75, 150 and 300 mM) (**Figure 2A**). We observed Cdt1 LLPS at or below physiological levels of salt (150 mM) but a loss of condensation at higher salt concentrations. These data demonstrate that electrostatic interactions are important for driving DNA-dependent phase separation by initiator proteins; however, it remained unclear whether specific Cdt1/DNA interactions, such as major- and minor-groove binding or ring-stacking interactions, were also necessary for LLPS. To address this question, we assessed whether other anionic polyelectrolytes of distinct chemical composition could drive condensation. Double-stranded (ds) and single-stranded DNA (ssDNA), dsRNA, heparin and poly-glutamate were tested (**Figure 2B**). Depletion assay results demonstrate that all polyanions tested showed an equal propensity to induce Cdt1 phase separation, strongly supporting the notion that non-specific electrostatic interactions are the primary driving force underlying DNA-induced initiator phase separation.

**Figure 2:**
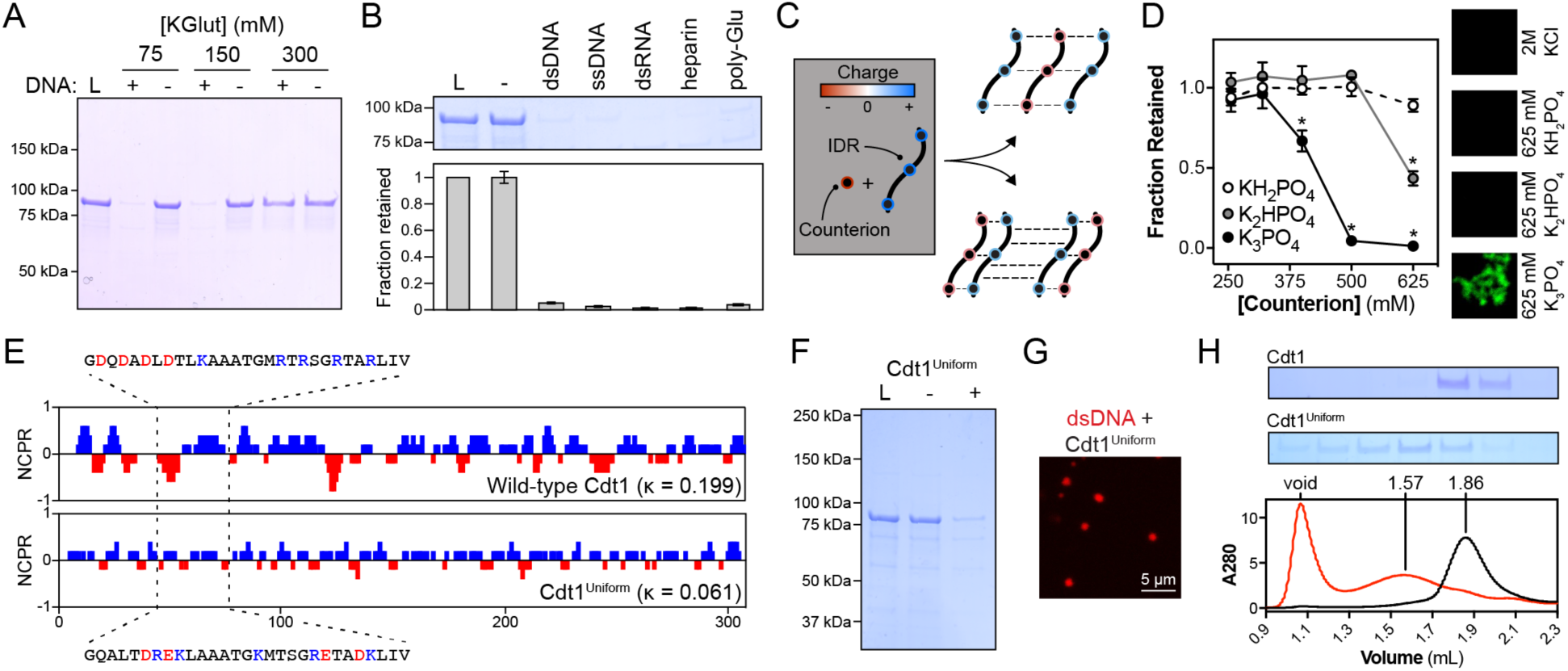
DNA contributes adhesive interactions to Cdt1 LLPS by acting as a counterion bridge. A) DNA-induced phase separation was assessed by the depletion assay in buffer containing 75, 150, or 300 mM potassium glutamate (KGlut). Cdt1 LLPS was observed at 75 and 150 mM KGlut, but not at 300 mM KGlut. B) Multiple anionic polyelectrolytes were assessed for their ability to induce Cdt1 phase separation by the depletion assay. “L” indicates a load control and “-” is in the absence of any anion polymer. dsDNA, ssDNA, dsRNA, heparin and poly-Glutamate (poly- Glu) all effectively induced Cdt1 LLPS. C) Two models explaining the ability of DNA to induce Cdt1 phase separation. (Top) DNA (black/red) functions as a counterion bridge to facilitate inter- IDR (blue/red) interactions. (Bottom) DNA neutralizes inter-IDR electrostatic repulsion to drive self-assembly. D) Monovalent (KH_2_PO_4_) and multivalent (K_2_HPO_4_ and K_3_PO_4_) phosphate counterions were assessed for the ability to induce Cdt1 phase separation by the depletion assay and fluorescence microscopy. For the line plot (left), the fraction Cdt1 retained in the depletion assay was quantitated and plotted against counterion concentration (mM). For the microscopy experiment (right), 5 µM eGFP-Cdt1 was combined with 2 M KCl, 625 mM KH_2_PO_4_, 625 mM K_2_HPO_4_ or 625 mM K_3_PO_4_ (top to bottom) and assessed for droplet formation by confocal fluorescence microscopy. Concentrated and highly networked species were observed only in the presence of K_3_PO_4_. E) Plot of the Net Charge Per Residue (NCPR) over a five-residue sliding window for Cdt1 and the charged residue variant, Cdt1^Uniform^. F) Depletion assay results for DNA- induced phase separation of Cdt1^Uniform^. “L” is a load control, “-“ is in the absence of DNA, and “+” is in the presence of DNA. G) Confocal fluorescence microscopy images of condensates formed with 5 µM Cy5-dsDNA and 5 µM Cdt1^Uniform^. H) Analytical size exclusion chromatography analysis of Cdt1 and Cdt1^Uniform^. Cdt1^Uniform^ adopts a more extended conformation in solution as evidenced by a lower retention volume and co-purifies with nucleic acid (260/280 = 0.56 and 0.79 for Cdt1 and Cdt1^Uniform^, respectively). “*” = p < 0.05.

In principle, two mechanisms could account for the dependence of Cdt1 LLPS on the presence of a counterion (**Figure 2C**). First, a counterion may function as an intermolecular bridge to contribute adhesive forces that help promote initiator LLPS. Alternatively, electrostatic repulsion between the positive charges on the Cdt1 IDR might prevent the spontaneous formation of inter-IDR interactions and LLPS, and a counterion could help neutralize this repulsive force. To distinguish between these mechanisms, we assayed phase separation by Cdt1 in the presence of mono- and multivalent counterions, with net charge ranging from -1 (KH_2_PO_4_) to -3 (K_2_HPO_4_) (**Figure 2D**). We reasoned that if counterions simply reduce electrostatic repulsion, then the titration of a monovalent salt would be sufficient to shield the IDR’s positive charge and drive phase separation. However, if counterions provide an intermolecular “bridge” between basic residues within the Cdt1 IDR, then multivalent salts would be required to drive phase separation. Using the depletion assay we found that monobasic potassium phosphate was unable to induce phase separation of Cdt1 up to the highest concentration tested (625 mM). Conversely, we observed significant depletion of Cdt1 at the highest concentration of dibasic potassium phosphate (625 mM), as well as for the three highest concentrations of tribasic potassium phosphate (400, 500 and 625 mM). eGFP-tagged Cdt1 (eGFP-Cdt1) and fluorescence microscopy were used to confirm the depletion assay results. In the presence of either 625 mM mono- or dibasic potassium phosphate, eGFP-Cdt1 appeared monodisperse; however, tribasic potassium phosphate induced formation of a concentrated eGFP-Cdt1 species. Interestingly, this species was morphologically distinct from the round droplets of Cdt1 that are observed in the presence of DNA and appeared more akin to aggregates than to droplets. These data indicate that the physiochemistry of a complexing polyanion can shape the properties of initiator condensates, consistent with previous observations for the complex coacervation of poly(proline-arginine) peptides (Boeynaems et al., 2019). Altogether, these data strongly indicate that counterions contribute an adhesive force to initiator self-assembly, bridging between the IDRs of different Cdt1 protomers.

In addition to overall net charge, recent theoretical and experimental work has demonstrated that the patterning of charged residues within disordered domains can fine-tune phase separation propensity (Lin et al., 2018; Nott et al., 2015; Pak et al., 2016; Paloni et al., 2020). We assessed the distribution of charged residues within the Cdt1 IDR by calculating the net charge per residue (NCPR) over a 5-residue window (**Figure 2E**, top). Ionic residues within the Cdt1 IDR cluster into local regions with net positive and net negative charge. We predicted that if charge distribution were important for initiator phase separation, then charge patterning would be conserved across metazoan initiator IDRs. To test this idea, we performed a kappa value analysis of initiator IDRs. Kappa is a parameter between 0-1 that quantitatively describes the degree of mixing between oppositely charged residues within a sequence (well-mixed samples have a low kappa) and serves as a predictor of IDR conformational state (extended or collapsed) (Das and Pappu, 2013). Given the absence of sequence conservation across initiator IDRs (Cdt1, Orc1, and Cdc6), we were surprised to find that the kappa values for metazoan initiator IDRs are similar (range 0.20-0.29), as are other physiochemical parameters (fraction charged residues, **FCR**, range 0.22-0.32; isoelectric point, pI, range 9.4-11.1), properties that are not conserved in budding yeast initiators (**Table 2**). To directly test whether the distribution of charged residues within the Cdt1 IDR are important for phase separation, we produced a Cdt1 variant, Cdt1^Uniform^ (kappa = 0.06), where we uniformly distributed charged residues (K, R, D, and E) across the IDR sequence, thereby retaining the same composition and pI as wild-type protein. A plot of Cdt1^Uniform^ NCPR demonstrates the loss of regions with high or low local net charge (**Figure 2E**, bottom). Utilizing the depletion assay (**Figure 2F**) and fluorescence microscopy (**Figure 2G**), we found that Cdt1^Uniform^ effectively induced phase separation in the presence of DNA, demonstrating that initiator phase separation is not strictly dependent on a specific ordering or clustering of charged residues.

**Table 2:** Charged residue properties of metazoan and yeast initiator IDRs.

| Species | Protein | FCR | Kappa | pI |
| --- | --- | --- | --- | --- |
| *Fruity fly* | Orc1 | 0.30 | 0.23 | 10.2 |
|  | Cdc6 | 0.31 | 0.25 | 9.4 |
|  | Cdt1 | 0.30 | 0.20 | 10.1 |
| *Human* | Orc1 | 0.31 | 0.21 | 10.6 |
|  | Cdc6 | 0.25 | 0.21 | 10.6 |
|  | Cdt1 | 0.28 | 0.26 | 10.6 |
| *Frog* | Orc1 | 0.32 | 0.29 | 9.7 |
|  | Cdc6 | 0.25 | 0.21 | 10.8 |
|  | Cdt1 | 0.30 | 0.20 | 10.1 |
| *Zebrafish* | Orc1 | 0.29 | 0.30 | 9.8 |
|  | Cdc6 | 0.22 | 0.20 | 11.1 |
|  | Cdt1 | 0.29 | 0.20 | 9.9 |
| *Budding yeast* | Orc1 | 0.48 | 0.43 | 4.7 |
|  | Cdc6 | 0.32 | 0.59 | 6.3 |
|  | Cdt1 | na | na | na |
| *Fission yeast* | Orc1 | 0.32 | 0.37 | 10.6 |
|  | Cdc6 | 0.21 | 0.21 | 10.2 |
|  | Cdt1 | na | na | na |

The dispensability of charged residue patterning for phase separation was initially surprising and prompted us to consider alternative roles for the observed charged residue patterning. A possible answer was gleamed from observations made during the purification of Cdt1^Uniform^. All proteins purified for this study were subject to a final polishing step over a size exclusion column, and it is at this step that we noticed a substantial difference in the elution profile of Cdt1 versus Cdt1^Uniform^. Wild-type Cdt1 eluted from a sizing column at a volume consistent with its physical parameters (size and globularity) and had no contaminating nucleic acid (A260/A280 = 0.56, 100% protein by mass). Conversely, Cdt1^Uniform^ eluted considerably earlier in a much broader peak, indicating that it possesses a more extended conformation compared to the native protein, and it co-eluted with a small fraction of nucleic acid (A260/A280 = 0.79, or 98% protein by mass) (**Figure 2H**). Together, these results indicate that the distribution of charged residues within the Cdt1 IDR encodes some type of higher-order structural or conformational information that also regulates nucleic-acid binding capacity.

### Intermolecular IDR-IDR interactions can drive LLPS in the presence of crowding reagent

A parsimonious mechanism for Cdt1 LLPS is one that relies exclusively on adhesive electrostatic interactions between the initiators and DNA (**Figure 3A**, top). However, it remained to be determined whether direct, inter-IDR contacts might provide an additional set of cohesive interactions that also promote LLPS (**Figure 3A**, bottom). To test this idea, we asked whether Cdt1 could form condensates in the absence of DNA (or any other appropriate counterion) by the addition of a crowding reagent (**Figure 3B**). Using the depletion assay, we titrated PEG-3350 (from 0-12.5% w/v in 2.5% increments) with Cdt1 and assayed protein depletion from the supernatant. We observed a statistically significant reduction of Cdt1 from the supernatant starting at the lowest concentration of PEG-3350 (2.5%) that progressed as PEG concentrations were increased to a near-complete depletion of Cdt1 from the supernatant at the highest value tested (12.5% PEG-3350). To confirm that the PEG-induced loss of protein in the depletion assay represented phase separation (as opposed to precipitation), fluorescence microscopy was used to visualize mixtures of eGFP-Cdt1 and PEG-3350. Mixtures of Cdt1 and 4% PEG-3350 were sufficient to induce clear droplet formation (**Figure 3C**, bottom panel). These data show that, in addition to poly-anions such as DNA, intermolecular Cdt1-Cdt1 interactions provide a driving force for LLPS.

**Figure 3:**
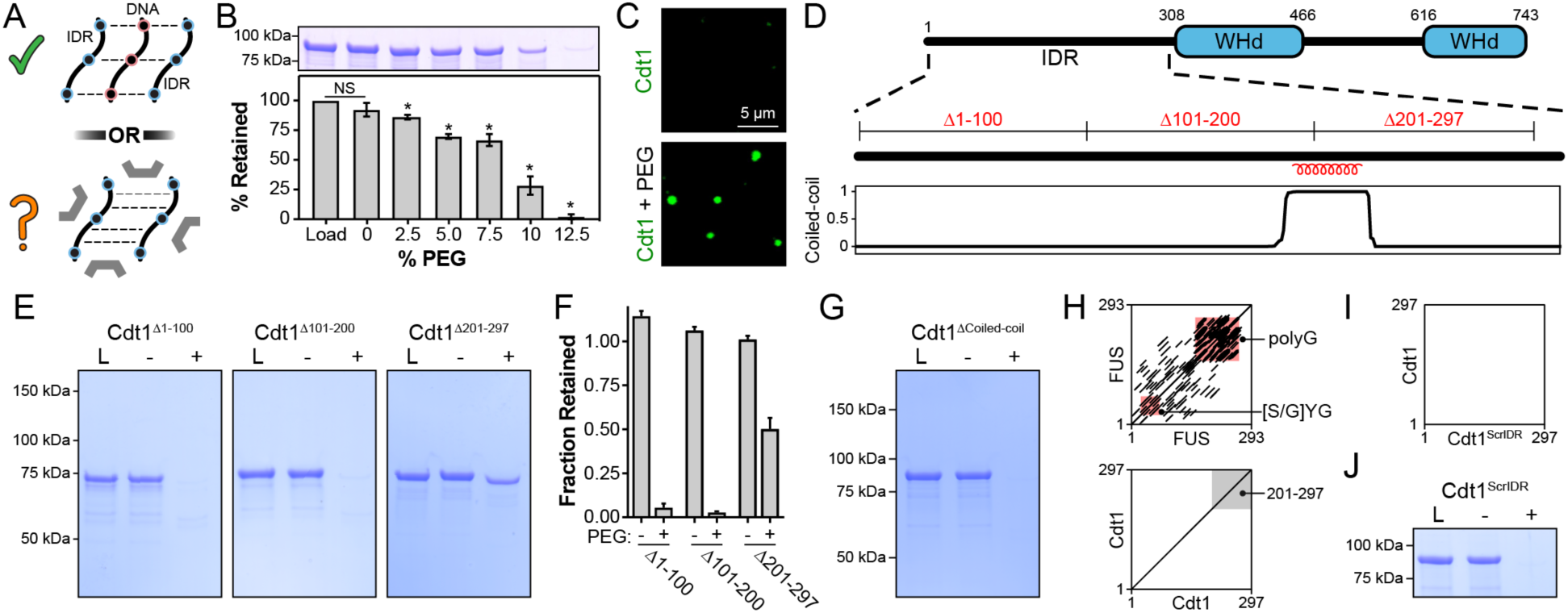
Inter-IDR cohesive interactions drive Cdt1 phase separation in the absence of DNA. A) (Top) DNA (black/red) can induce Cdt1 (black/blue) phase separation by acting as a counterion bridge. (Bottom) To what extent homotypic inter-IDR interactions affect phase separation was unknown. B) Depletion assay to assess phase separation of 8 µM Cdt1 in the presence of increasing concentrations of PEG-3350 (0-12% w/v). C) Phase separation of 5 µM eGFP-labeled Cdt1 was assessed by confocal fluorescence microscopy in the absence (top) and presence (bottom) of 4% PEG-3350. D) Schematic representation of the domain architecture of *D. melanogaster* Cdt1 (top) with coiled-coil domain prediction for the Cdt1 N-terminal IDR (bottom). The position of the three approximately 100 amino acid deletions used in Figure 3E are indicated. E) Cdt1^Δ1-100^ (left), Cdt1^Δ101-200^ (middle), and Cdt1^Δ201-297^ (right) were assayed for the ability to undergo PEG-induced phase separation by the depletion assay. “L” is a load control, “-“ is in the absence of PEG, and “+” is in the presence of PEG. F) Quantitation of depletion assay results presented in Figure 3E. G) The depletion assay was used to assess phase separation of a Cdt1 variant lacking the coiled-coil domain (Cdt1^ΔCoiled-coil^). “L” is a load control, “-“ is in the absence of PEG, and “+” is in the presence of PEG. H) Dotplot analysis (EMBOSS Dotmatcher, (Madeira et al., 2019)) of repetitive elements in the human FUS IDR (top) and the Cdt1 N-terminal IDR (bottom). The region representing Cdt1 amino acids 201-297 are highlighted with a grey background. The main diagonal represents sequence homology with itself and diagonals off the main diagonal represent repetitive motifs. Many repetitive motifs are apparent in FUS but are lacking in Cdt1. I) Dotplot analysis of the Cdt1^ScrIDR^, a Cdt1 variant with N-terminal IDR residues randomly scrambled, versus the wild-type Cdt1 IDR. J) Depletion assay to assess phase separation of Cdt1^ScrIDR^. “L” is a load control, “-” is in the absence of PEG, and “+” is in the presence of PEG.

The ability of Cdt1 to self-associate prompted us to look for specific motifs within its IDR that could promote such interactions. As a first step, three different 100-amino acid segments were removed from the Cdt1 IDR (**Figure 3D**) and assessed for how the deletions impacted PEG-induced Cdt1 phase separation (**Figure 3E**). Deletion of IDR residues 1-100 (Cdt1^Δ1-100^) or 101-200 (Cdt1^Δ101-200^) had no discernible effect on Cdt1’s propensity to phase separate in the presence of PEG. However, we observed a modest reduction in Cdt1 partitioning (∼50%) into the condensed phase when residues 201-297 (Cdt1^Δ201-297^) were deleted (**Figure 3F**). Although there is a predicted coiled-coil domain within the final third of the Cdt1 IDR (residues 196-223) that was suggestive of a possible self-association motif (**Figure 3D**, bottom), the deletion of this region had no impact on Cdt1’s condensation properties (**Figure 3G**).

Protein multivalency is a key property of proteins that undergo LLPS and frequently can arise from disordered sequences that contain short repetitive interaction motifs (Banani et al., 2017). In the absence of obvious structural features to explain Cdt1 self-association, we attempted to identify repetitive motifs within the Cdt1 IDR – particularly within the 201-297 region – that might explain Cdt1 self-assembly. A prototypic example of repetitive IDR elements are those contained within human FUS, whose IDR contains multiple [S/G]YG motifs that are important for self-association (Kato et al., 2012; Luo et al., 2018). These repeats, in addition to the protein’s poly-glycine sequences, can be visualized diagrammatically by a Dotplot (Madeira et al., 2019) (**Figure 3H**, top). In this plot, a sequence is plotted against itself and a dot is placed at each point where a repetitive match of a specific length and threshold is found. We used this approach to search for repetitive sequence elements within the Cdt1 IDR. However, in stark contrast to FUS and many other condensate forming proteins (**Figure 3-figure supplement 1A-F**), we observed no repetitive motif(s) within the Cdt1 IDR (**Figure 3H**, bottom). This result led us to hypothesize that Cdt1 lacks repetitive sequences that would help promote self-association, and that it is the IDR amino acid composition alone that contains the necessary components to promote inter-protomer interactions and phase separation. To test this idea, we synthesized and produced a Cdt1 variant, Cdt1^ScrIDR^, where the order of the N-terminal IDR amino acids (residues 1-297) were randomly scrambled and then fused back to the wild-type C-terminal portion of the protein (residues 298-743). The depletion assay was then used to ask whether this protein was still capable of phase separation. Surprisingly, Cdt1^ScrIDR^ showed robust phase separation propensity in the presence of a crowding agent, demonstrating that Cdt1^ScrIDR^ retains the self-associative capacity observed for wild-type Cdt1 (**Figure 3J**). Notably, Cdt1^ScrIDR^ also formed condensates in the presence of DNA (**Figure 3-figure supplement 2A**) and these were able to recruit eGFP- Cdt1, although the efficiency of recruitment was lower than that observed with wild-type Cdt1 condensates (**Figure 3-figure supplement 2B-C**). Collectively, these data demonstrate that it is the overall amino acid composition of initiator IDR sequences, not sequence order, that is essential for Cdt1’s self-associative capacity and that composition alone can promote heteromeric inter-IDR interactions.

### Branched hydrophobics are disproportionately enriched in the Cdt1 IDR and drive LLPS

The Cdt1 IDR contains alternating blocks of net-positive and net-negatively charged residues (**Figure 2E**). We reasoned that these interspersed blocks of opposite charge might facilitate inter-IDR interactions in the presence of a crowding reagent to drive phase separation; other condensate-forming proteins, such as *C. elegans* Laf1 (Elbaum-Garfinkle et al., 2015) and Ddx4 (Nott et al., 2015), have been shown to phase separate in such a manner. To test this idea, we compared the salt-sensitivity of PEG-induced Cdt1 LLPS to DNA-induced condensation (**Figure 4A**). At 75 mM and 150 mM KGlutamate, both DNA and PEG were comparable in their ability to induce Cdt1 phase separation. By contrast, salt concentrations above 150 mM KGlutamate abolished DNA-induced Cdt1 phase-separation but had no effect on PEG-induced LLPS. This result demonstrates that charge-charge interactions are not a major contributor to the inter-IDR interactions that drive PEG-induced phase separation.

**Figure 4:**
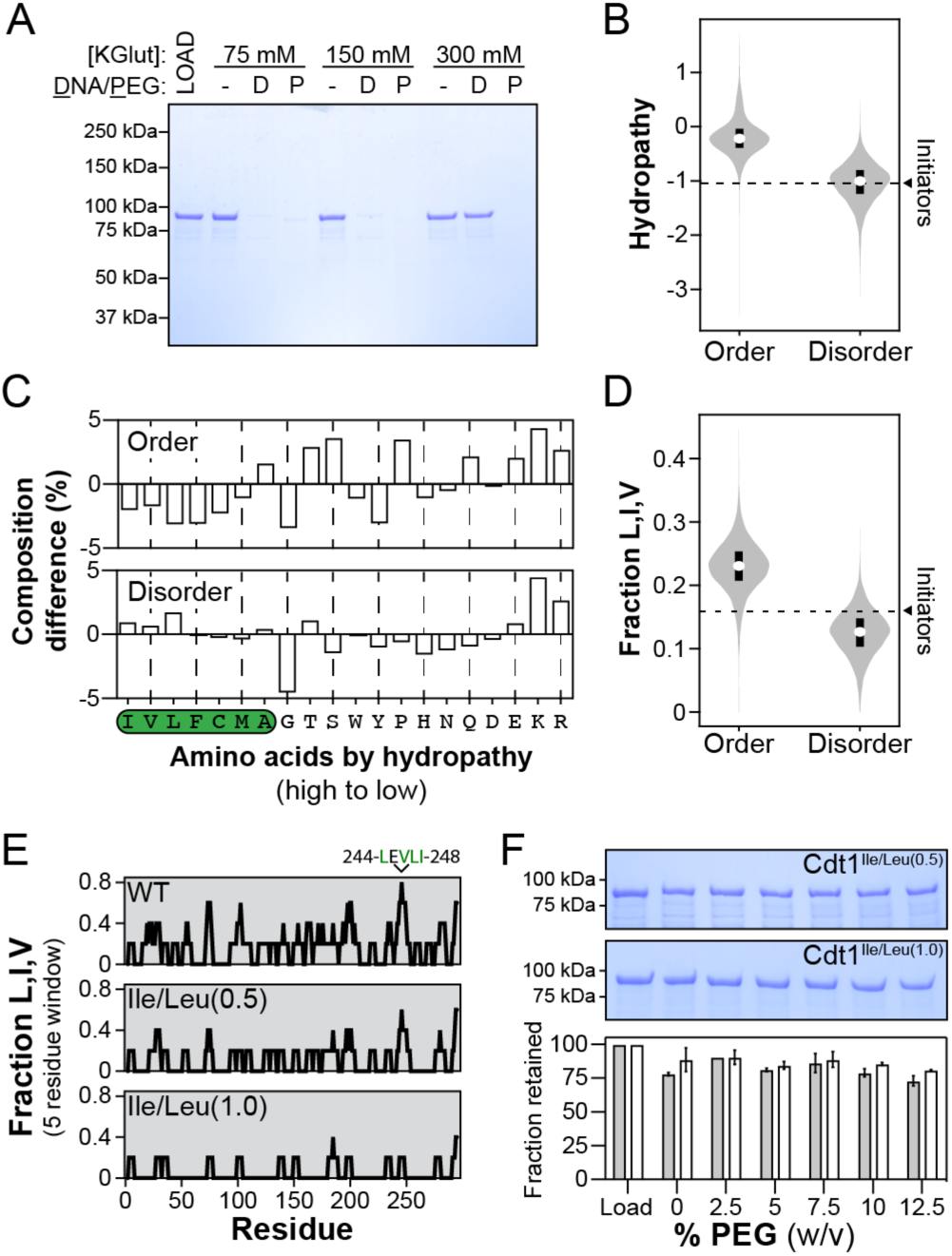
Branched hydrophobic residues underlie Cdt1 homotypic inter-IDR interactions. A) The depletion assay was used to assess the salt-sensitivity of PEG-induced phase separation side-by-side with DNA-induced phase separation. “L” is a load control, “-” is in the absence of DNA and PEG, “D” indicates the addition of DNA and “P” indicates the addition of PEG-3350. Cdt1 phase separation was assessed at 75, 150 and 300 mM KGlutamate (“KGlut”). B) Comparison of initiator IDR hydropathy (dashed line indicates the average hydropathy of the Orc1, Cdc6 and Cdt1 IDRs) to the hydropathy of predicted ordered domains (“Order”) and disordered domains (“Disorder”) proteome wide. C) Percent difference in initiator IDR amino acid composition relative to all predicted ordered (top) and disordered (bottom) domains proteome wide. The amino acids are ordered from high to low (left to right) hydropathy and those with a positive hydropathy value are indicated by a green background. D) Comparison of initiator IDR fraction Leu, Ile and Val (dashed line indicates the average fraction L, I, V of the Orc1, Cdc6 and Cdt1 IDRs) to the fraction L, I, V of all predicted ordered domains (“Order”) and disordered domains (“Disorder”) proteome wide. E) To assess the distribution of branched hydrophobic residues, the fraction L, I, V was calculated over a 5-residue sliding window for the Cdt1 IDR (top). Two mutants were made with either half (Cdt1^Ile/Leu(0.5)^) or all (Cdt1^Ile/Leu(1.0)^) of Leu and Ile residues mutated to Ala, and the distribution of L, I, V was calculated for each (middle and bottom, respectively). F) The depletion assay was used to assess the phase separation capacity of Cdt1^Ile/Leu(0.5)^ (top gel) and Cdt1^Ile/Leu(1.0)^ (bottom gel) in the presence of increasing concentrations of PEG-3350 (0-12% w/v). The fraction of protein retained at each concentration of PEG-3350 was quantitated.

We next set out to test whether Cdt1 self-assembly might instead be driven by the hydrophobic effect. We first asked whether initiator IDRs may be uniquely enriched in hydrophobic residues. However, when we compared the hydropathy of initiator IDRs to all ordered and disordered segments longer than 100 amino acids within the *D. melanogaster* proteome (14,002 ordered domains and 4957 disordered domains), we found that initiator IDRs were closer to the hydropathy of the median IDR (average of -1.04 for Orc1, Cdc6, and Cdt1) than to either the lower (-1.24) or upper (-0.80) quartile of the data (**Figure 4B**). This result led us to take a more unbiased approach and compare initiator IDR sequence composition (fraction average of each amino acid across Orc1, Cdc6, and Cdt1 IDRs) to the average composition of all ordered and disordered domains longer than 100 amino acids (**Figure 4C**). We found that initiator IDRs are more highly enriched in both the three most hydrophobic residues (isoleucine, valine and leucine) and the three least hydrophobic residues (glutamate, lysine and arginine) compared to other disordered domains. We also compared the total fraction of Leu, Ile, and Val (fraction-L,I,V) residues in the initiators to all ordered and disordered domains proteome wide. We found that the initiator fraction-L,I,V is within the top 25% of all disordered domains (**Figure 4D**).

The unique enrichment of branched hydrophobic residues within initiator IDRs prompted us to test whether these amino acid types promote adhesive inter-IDR interactions that drive Cdt1 self-association. The fraction-L,I,V in the Cdt1 IDR was first calculated over a five-residue sliding window to determine whether hydrophobic residues were uniformly distributed throughout the sequence or clustered into discrete motifs targetable by mutagenesis (**Figure 4E**). Overall, the Cdt1 IDR bears an average fraction-L,I,V of 0.19 – or approximately one Leu, Ile, or Val for every 5 residues – and the hydrophobic residues are generally well-distributed throughout the sequence. There are five 5-residue windows where the fraction-L,I,V reaches 0.6, and each third of the IDR (residues 1-100, 101-200, 201-297) contains at least one of these regions. A fraction- L,I,V = 0.8 occurs only once, between residues 244-248. Interestingly, the C-terminal third of the Cdt1 IDR (residues 201-297) contributes more strongly to PEG-induced LLPS than either the first (1-100) or middle third (101-200) of the sequence (**Figure 3E**), suggesting that the 244-248 sequence may underlie the greater cohesiveness of this region. A higher-precision analysis of Cdt1’s hydrophobic residue distribution shows that 67% of Leu, Ile, and Val residues reside in isolation, 22% are immediately adjacent to another Leu, Ile, or Val residue, and 11% are within tri-peptides composed of the three amino acids.

Because sequence analysis shows that hydrophobic residues are broadly distributed across the length of the Cdt1 IDR, we were interested to test whether these amino acids contribute to PEG-induced LLPS in an accumulative manner. Two Cdt1 IDR mutants were constructed to probe this issue, one with half of all Ile and Leu residues mutated to alanine (Cdt1^Ile/Leu(0.5)^) and another with all Ile and Leu residues mutated (Cdt1^Ile/Leu(1.0)^). Ile and Leu were chosen as targets, as these are the two most highly enriched hydrophobic residues in initiator IDRs compared to disordered sequences across the proteome. To assess phase separation potential, PEG was titrated (from 0-12.5% in 2.5% increments) against each mutant and their disappearance monitored from the supernatant by depletion assay (**Figure 4F**). The phase separation propensity of both variants was significantly disrupted. At the highest concentration of PEG (12.5%), we observed only a modest loss of Cdt1^Ile/Leu(0.5)^ and Cdt1^Ile/Leu(1.0)^ (27% and 19%, respectively) from the supernatant, compared to nearly a 100% loss for wild-type Cdt1 (**Figure 3B**). These data establish that, relative to the average disordered domain, initiator IDRs are enriched in branched hydrophobic residues and these amino acid types mediate adhesive inter-IDR interactions that underlie phase separation by Cdt1 under conditions of molecular crowding.

### Inter-IDR hydrophobic interactions are necessary for DNA-induced LLPS

Our results have shown that both DNA-bridging interactions (mediated by electrostatics) and inter-IDR interactions (mediated by hydrophobic residues) are important for phase separation by Cdt1. However, the extent to which these two types of interactions synergize remained unclear. We therefore set out to determine whether inter-IDR interactions promote Cdt1 phase separation in the presence of DNA. During the purification of Cdt1^Ile/Leu(0.5)^ and Cdt1^Ile/Leu(1.0)^ we noticed that both proteins showed a normal affinity for heparin (**Figure 5 – figure supplement 1**), indicating that the mutants might bind DNA with wild-type-like affinity. This assumption was confirmed by electrophoretic mobility shift assay, which showed that the DNA-binding affinity of Cdt1^Ile/Leu(0.5)^ (K_d, apparent_ = 67 nM) and Cdt1^Ile/Leu(1.0)^ (K_d, apparent_ = 106 nM) is similar to that of the wild-type protein (K_d, apparent_ = 69 nM) (**Figure 5A**). These data establish Cdt1^Ile/Leu(0.5)^ and Cdt1^Ile/Leu(1.0)^ as separation- of-function mutants that are capable of binding DNA (**Figure 5A**) but that are deficient in their ability to participate in inter-IDR interactions (**Figure 4**).

**Figure 5:**
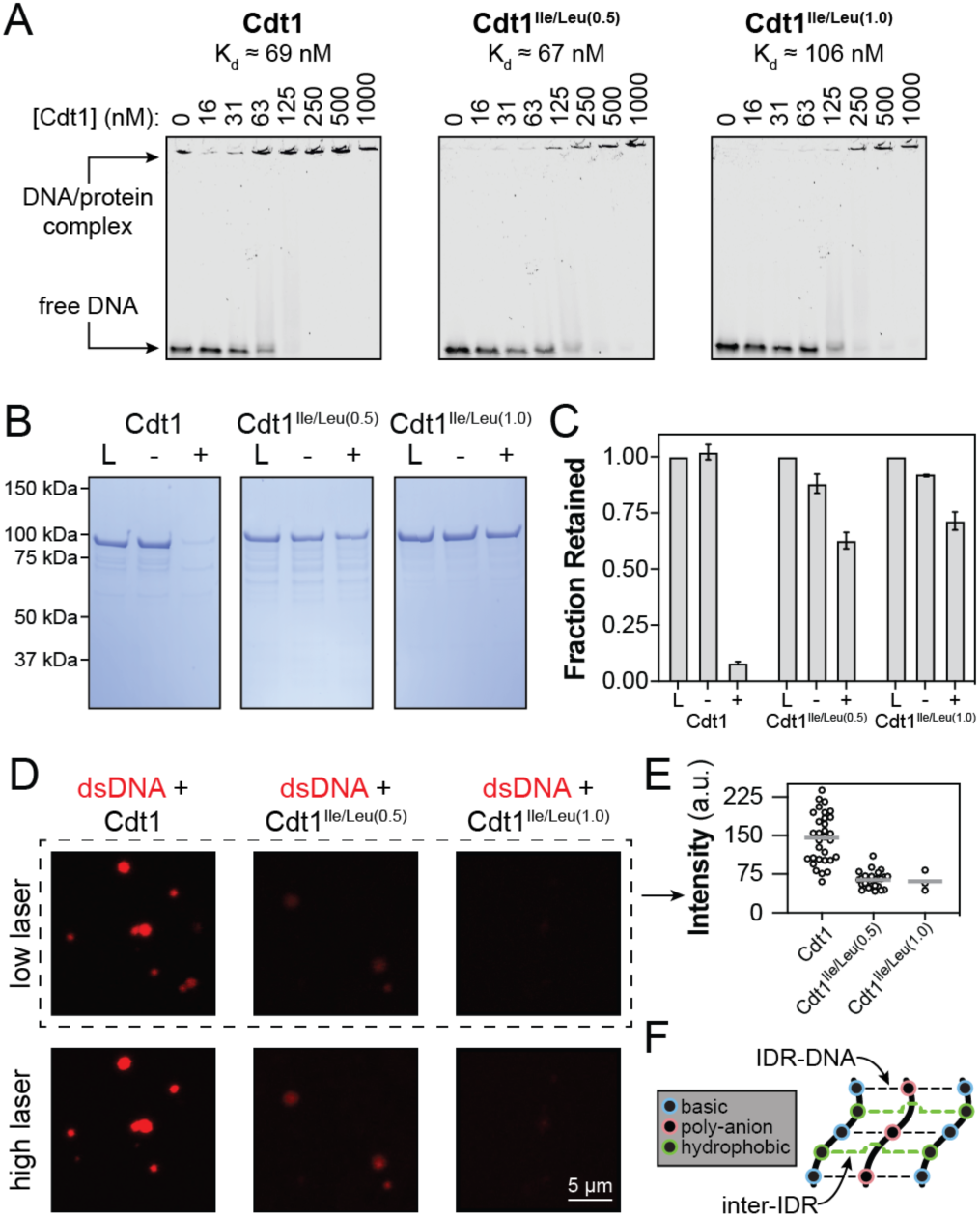
Homotypic inter-IDR interactions underlie DNA-induced initiator phase separation. A) An electrophoretic mobility shift assay (EMSA) was used to assess the binding affinity of Cdt1 (left), Cdt1^Ile/Leu(0.5)^ (middle) and Cdt1^Ile/Leu(1.0)^ (right) for dsDNA. Binding was assessed with 10 nM IRDye800-labeled dsDNA and a titration of Cdt1 from 16 to 1000 nM. Each variant showed comparable affinity to wild-type Cdt1. B) Depletion assay results assessing DNA-induced phase separation of Cdt1 (left), Cdt1^Ile/Leu(0.5)^ (middle) and Cdt1^Ile/Leu(1.0)^ (right) . “L” is a load control, “-” is in the absence of DNA, and “+” is in the presence of DNA. C) Quantitation of the depletion assay results presented in Figure 5B. D) Confocal fluorescence microscopy was used to assess DNA-induced phase separation by combining 5 µM Cdt1 (left) and the Cdt1 variants Cdt1^Ile/Leu(0.5)^ (middle) and Cdt1^Ile/Leu(1.0)^ (right) with 5 µM Cy5-dsDNA. Images were taken under low (top row) and high (bottom row) laser power to assess for weakly fluorescent condensates. E) Micrographs of Cy5-dsDNA/Cdt1 droplets (Figure 5D) were thresholded to identify condensates and the maximum Cy5-dsDNA signal observed within each droplet was measured. The data include observations from three separate micrographs for each Cdt1 variant. F) Initiator IDR phase separation relies on synergistic heteromeric IDR-DNA and homomeric IDR-IDR interactions. Positively charged residues within the IDR (blue) contribute to self-assembly by interacting with DNA (red) which can bridge inter-IDR interactions (black dotted lines). Branched hydrophobic residues (green) mediate direct homotypic inter-IDR interactions through the hydrophobic effect (green dotted lines).

We next utilized these mutants to ask whether DNA bridging by itself is sufficient to drive Cdt1 LLPS, or whether inter-IDR cohesive interactions are also required. We first used the depletion assay to compare DNA-induced phase separation of Cdt1^Ile/Leu(0.5)^ and Cdt1^Ile/Leu(1.0)^ vs. the wild-type protein (**Figure 5B**). While wild-type Cdt1 showed near complete partitioning into the condensed phase (8% protein retention in the supernatant), 63% of Cdt1^Ile/Leu(0.5)^ and 73% of Cdt1^Ile/Leu(1.0)^ remained in the supernatant, demonstrating a marked reduction in DNA-induced phase separation (**Figure 5C**). To confirm these results and to assess the nature of the condensed phase that forms with mutant proteins, we utilized fluorescence microscopy to visualize droplets in the presence of Cy5-dsDNA (**Figure 5D**). Compared to wild-type Cdt1, which robustly formed droplets (left set of images), only a few, weakly fluorescent droplets were seen in solutions containing Cy5-dsDNA and Cdt1^Ile/Leu(0.5)^ (middle set of images), and even fewer droplets of weaker intensity were observed for Cdt1^Ile/Leu(1.0)^ (right set of images). We imaged the same fields of view at higher laser intensity to ensure that droplets of very weak intensity did not go undetected (bottom set of images). Quantification across three replicate images of Cy5 signal intensity within droplets revealed that both mutant constructs displayed a > 50% loss in mean signal intensity compared to wild-type Cdt1 (we anticipate that the Cdt1 concentration within these droplets is also dramatically reduced, which would be consistent with our depletion assay results, **Figure 5B**). These data demonstrate a striking reduction in the ability of Cdt1^Ile/Leu(0.5)^ and Cdt1^Ile/Leu(1.0)^ to undergo DNA-induced phase separation, despite possessing normal DNA binding activity, and suggest that cohesive inter-IDR interactions play a uniquely critical role in driving the phase separation propensity of Cdt1.

## DISCUSSION

In previous work, we discovered that several factors used to initiate DNA replication in metazoans – in particular the Orc1 subunit of ORC, as well as Cdc6 and Cdt1 – possess an N-terminal IDR that promotes protein phase separation upon binding DNA (Parker et al., 2019). When these initiator proteins are combined and DNA is added, they form co-mingled condensates that exclude non-cognate phase-separating proteins and are active for the ATP-dependent recruitment of the Mcm2-7 helicase. The biophysical mechanisms underlying LLPS by replication initiators have remained unknown. Here, we have used *Drosophila* Cdt1 as a model system to dissect the molecular basis for DNA-dependent phase separation by initiation factors. Our studies show that this phase separation mechanism has several unique features compared to other condensate-forming proteins characterized to date.

We find that two distinct types of intermolecular interactions synergize with each other to drive initiator LLPS, and that one of these dominates over the other. Interactions are shown to form between Cdt1 and DNA that are primarily non-specific and electrostatic in nature, and in which DNA serves as a counterion bridge between different Cdt1 molecules (**Figure 2** and **Figure 5F**, black dotted lines). Inter-IDR interactions that are salt-insensitive and primarily hydrophobic in nature are also identified (**Figure 4** and **Figure 5F**, green dotted lines). Importantly, we find that inter-IDR interactions can drive Cdt1 LLPS in the absence of DNA (**Figure 3**), but that when these interactions are abolished, DNA is insufficient to induce phase separation (**Figure 5**). Thus, inter-IDR interactions provide the primary force behind initiator LLPS, with Cdt1-DNA interactions contributing an additional adhesive force.

In the absence of a crowding reagent, DNA is required to induce initiator phase separation at physiological salt and protein concentrations (Parker et al., 2019). This dependency suggests that DNA serves as an essential nucleating determinant of where Cdt1 can form condensates, ensuring that initiator LLPS is restricted to a chromatin context. Interestingly, other anionic biopolymers, such as RNA and heparin, are also capable of inducing phase separation by Cdt1. This type of scaffold ‘promiscuity’ has been observed in other systems that undergo LLPS, such as for the nucleolar protein fibrillarin (Feric et al., 2016). Given that the cytosol contains multiple anionic biopolymers, future work will need to answer how replication initiators are specifically targeted to chromatin, a question that is also relevant for RNA-dependent cellular bodies (Berry et al., 2015; Elbaum-Garfinkle et al., 2015; Feric et al., 2016; Guillén-Boixet et al., 2020; Mitrea et al., 2016; Molliex et al., 2015; Van Treeck et al., 2018). The selection of an appropriate nucleic acid scaffold across different phase-separating systems is likely to rely on both general mechanisms, such as the sequestration of alternative, inappropriate scaffolds by proteins with different or non-existent LLPS properties, and by system-specific mechanisms, such as a requirement for particular sequences or secondary structure (Langdon et al., 2018; Maharana et al., 2018).

A growing body of work has demonstrated the importance of aromatic residues in driving protein phase separation (Chiu et al., 2020; Chong et al., 2018; Lin et al., 2017; Nott et al., 2015; Pak et al., 2016; Qamar et al., 2018; Wang et al., 2018), along with a related role for π-π and π-cation interactions (Vernon et al., 2018). In parallel, many studies have shown that condensates formed through aromatic residue interactions are readily dissolved by 1,6-HD (**Table 1**). A distinguishing feature of Cdt1 LLPS is that it is both resistant to treatment with 1,6-HD and does not rely on aromatic residues within its IDR for phase separation (**Figure 1**). We have also shown that ORC and Cdc6 phase-separation is resistant to treatment with 1,6-HD (**Figure 1-figure supplement 1**), and while we have not explicitly investigated the role of aromatic residues within these proteins, we note that the fraction aromatic residues of Orc1 (0.019) and Cdc6 (0.012) is lower than that of Cdt1 (0.027), suggesting that aromaticity is dispensable for LLPS by replication initiation factors in general.

Instead of relying on aromatic-mediated π-π or π-cation interactions, our data show that interactions between initiator IDRs are instead facilitated by branched hydrophobic residues. Although hydrophobic sequences have been shown to be important for the phase-separation of other factors, such as the extracellular matrix protein elastin (Quiroz and Chilkoti, 2015; Reichheld et al., 2017; Yeo et al., 2011) and the Nephrin intracellular domain (NICD) (Pak et al., 2016), important mechanistic distinctions exist with respect to metazoan replication initiators. For example, both NICD and Cdt1 bear an identical fraction of charged residues (FCR = 31%) and rely on a counterion for phase separation. However, inter-IDR interactions within the NICD appear to rely primarily on sequence aromatics (11% aromatic content), with hydrophobic residues playing only a supporting role in NICD self-association (Pak et al., 2016). Elastin, like metazoan replication initiators, is relatively devoid of aromatic residues (3.9%) and contains a highly similar content of branched hydrophobic amino acids that underly phase separation (elastin and Cdt1 fraction-L/I/V = 0.21 and 0.19, respectively), but differs in that it readily forms condensates in the absence of a counterion. Elastin’s hydrophobic sequences are contained within highly repetitive “VPGVG” and “GLG” sequences, and scrambling or shuffling these motifs negatively impacts elastin phase-separation (Pepe et al., 2008, 2005; Toonkool et al., 2001). Conversely, initiator IDRs lack such repeats and (for Cdt1 at least) are insensitive to sequence order. We speculate that inter-IDR interactions in initiators are weakened compared to elastin due to the non-repetitive, distributed nature of their hydrophobic resides and that additional counterion-mediated adhesive interactions are required to promote Cdt1 self-assembly.

Understanding the distinctive biophysical underpinnings of the different phase-separating systems is essential for developing *in silico* approaches for predicting the cellular partitioning of a given IDR from primary sequence alone. Such efforts will ultimately require an understanding of both the specific and general features of condensates’ scaffolding components (e.g. repetitive motifs vs. amino acid composition). The present work highlights the particular importance of general sequence features. We find that the Cdt1 IDR can be fully scrambled without losing the ability to form condensates (**Figure 3J**) and that condensates formed from the scrambled variant can recruit wild-type Cdt1 (**Figure 3** and **Figure 3-figure supplement 2A-C**). Similarly, regions of the NICD can be scrambled without losing the ability to form cellular condensates (Pak et al., 2016) and the LCDs of certain transcription factors can be shuffled without affecting promoter targeting (Brodsky et al., 2020). These studies demonstrate the importance of sequence composition in not only driving phase separation but also in facilitating interactions with partner-proteins, and is consistent with recent work showing that proteins with similar functional annotation have disordered sequences with similar, evolutionary conserved molecular features (pI, composition, FCR, kappa, etc.) (Zarin et al., 2019). Our observations involving the co-condensation properties of Cdt1, ORC and Cdc6 – three proteins with IDRs of similar amino acid composition but no direct sequence homology (Parker et al., 2019) – provide experimental support for this “like-recruits-like” hypothesis. Our previous work also identified compositional homology across metazoan initiator IDRs and led us to predict that the capacity to phase separate was likely conserved across metazoan initiators (Parker et al., 2019). These predictions were recently confirmed for human Orc1 and Cdc6 with the demonstration of their co-condensation in the presence of DNA, work that further emphasizes the importance of sequence composition (Hossain et al., 2021).

Defining the molecular ‘grammar’ that encodes LLPS in metazoan initiators likewise has implications for understanding the nature and function of their co-assembly *in vivo*. Work in multiple model organisms has revealed a switch-like transition in initiator localization during anaphase, during which initiators begin to coat chromosomes (Baldinger and Gossen, 2009; Kara et al., 2015; Parker et al., 2019; Sonneville et al., 2012). We predict that initiator self-assembly underlies this behavior and produces a replication layer, a dense region of chromatin-bound initiators poised for pre-replication complex (Pre-RC) assembly, that drives the precipitous, genome-wide loading of the Mcm2-7 complex in late mitosis (Dimitrova et al., 2002; Méndez and Stillman, 2000; Okuno et al., 2001). This moment in the cell cycle is an opportune time to prepare for replication, as conflicts with the transcriptional machinery are minimized (Palozola et al., 2017) and the chromatin substrate is, relative to interphase, uniformly compacted (Ou et al., 2017). The identification of Cdt1 mutants that selectively block phase separation, but not DNA binding (**Figure 5**), affords an opportunity to directly investigate the physical nature of initiator assemblies *in vivo*. Beyond helicase loading and licensing, initiator LLPS might also serve to prime replisome assembly. Following helicase loading, Mcm2-7 is phosphorylated by the Dbf4-dependent kinase (DDK), a heterodimer of Cdc7 and Dbf4 (Deegan et al., 2016; Sheu and Stillman, 2010; Yeeles et al., 2015). Intriguingly, the *Drosophila* homolog of Dbf4, Chiffon, contains multiple IDRs, and a large internal IDR (residues 903-1209) that possesses striking compositional similarity to initiator-type IDRs (**Figure 5-figure supplement 2**).

The present work provides a detailed view of the DNA-dependent liquid-liquid phase separation mechanism for *Drosophila* Cdt1. Due to a high-level of IDR compositional homology and an ability to form co-mingled phases (Parker et al., 2019), the unique mechanism we describe for Cdt1 phase-separation likely extends to the other replication initiation factors, ORC and Cdc6. These studies set the stage for investigating the physiological significance of initiator self-assembly in replication licensing and for identifying other sequences in the proteome that possess IDRs capable of associating with chromatin and initiation factors alike.

## MATERIALS AND METHODS

### Protein production and purification

Wild-type and mutant Cdt1 coding sequences (**Table 3**) were cloned into the QB3 Macrolab vector 2CT to append a tobacco etch virus (TEV) protease-cleavable C-terminal hexa-histidine (His6)-maltose binding protein (MBP) tag. To produce eGFP-tagged Cdt1, the wild-type Cdt1 coding sequence was cloned into QB3 Macrolab vector 1GFP to append an N-terminal His6-eGFP tag. All Cdt1 variants were produced from Rosetta 2(DE3) pLysS cells. Cells were grown in 2xYT broth at 37°C to an OD600 = 0.8 and after a 15 min incubation in an ice bath induced with 1 mM isopropyl β-d-thiogalactoside (IPTG). After growth overnight at 16°C, cells were harvested by centrifugation and cell pellets frozen at -80°C until further processing.

**Table 3:**
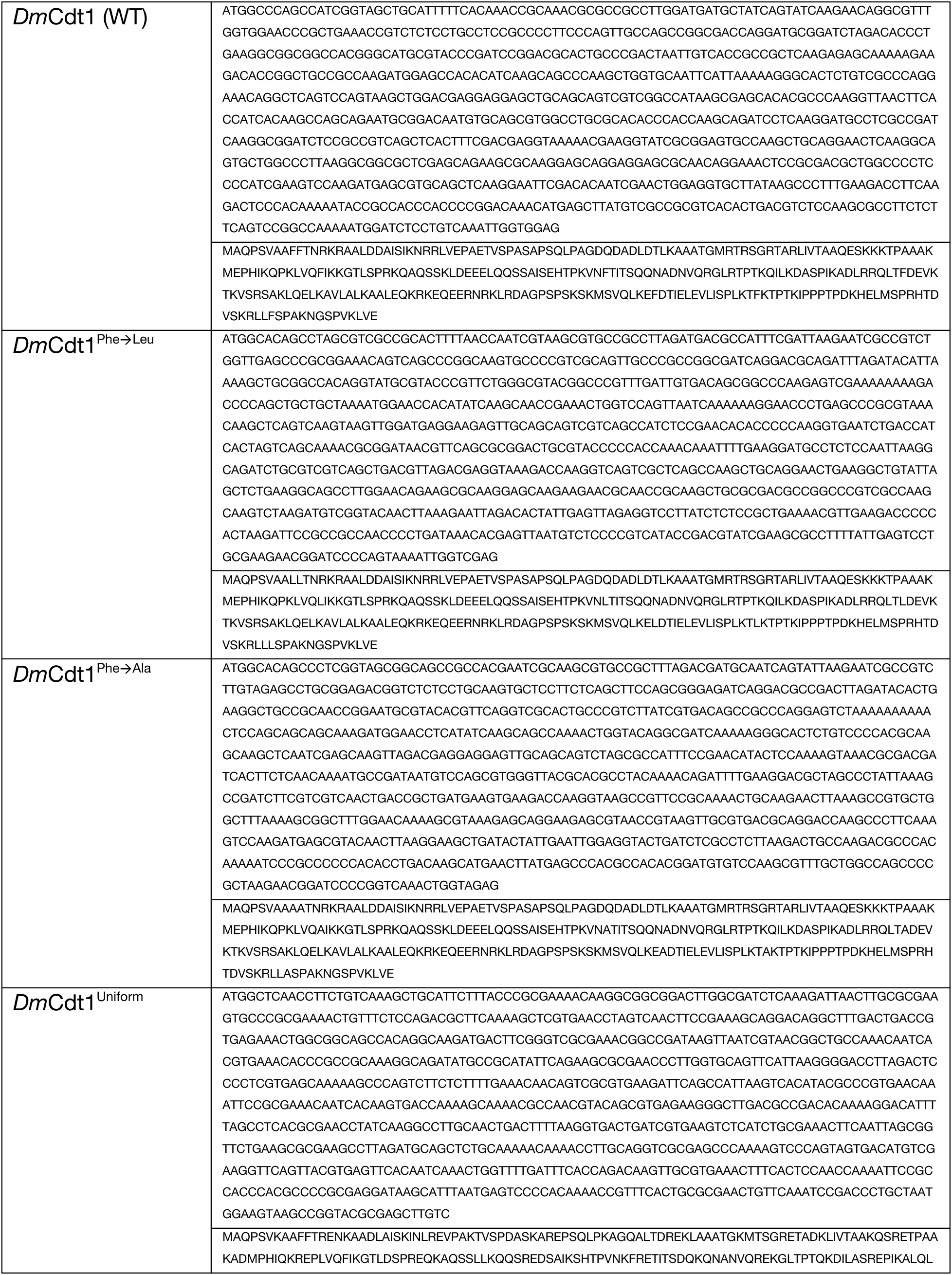
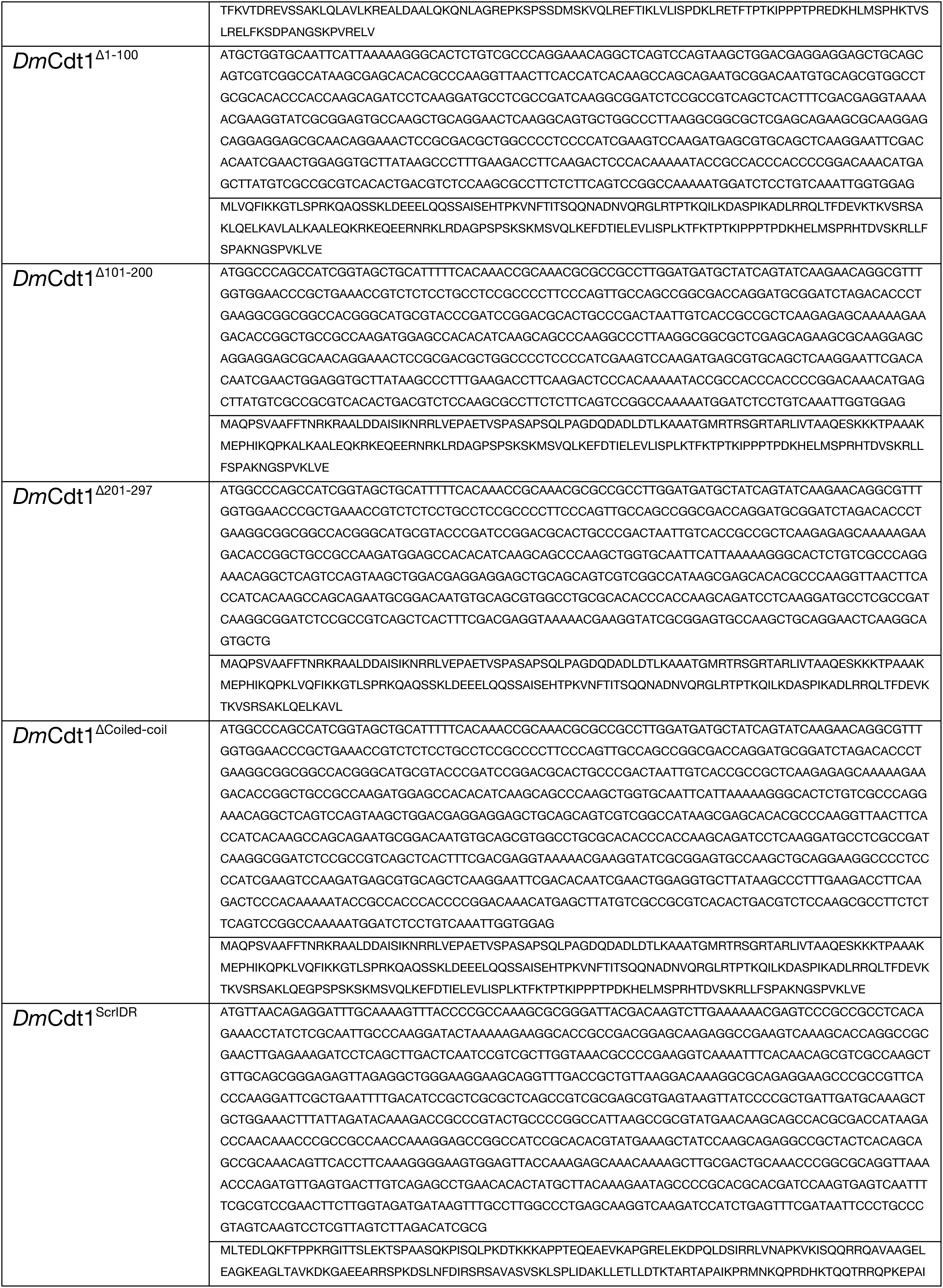
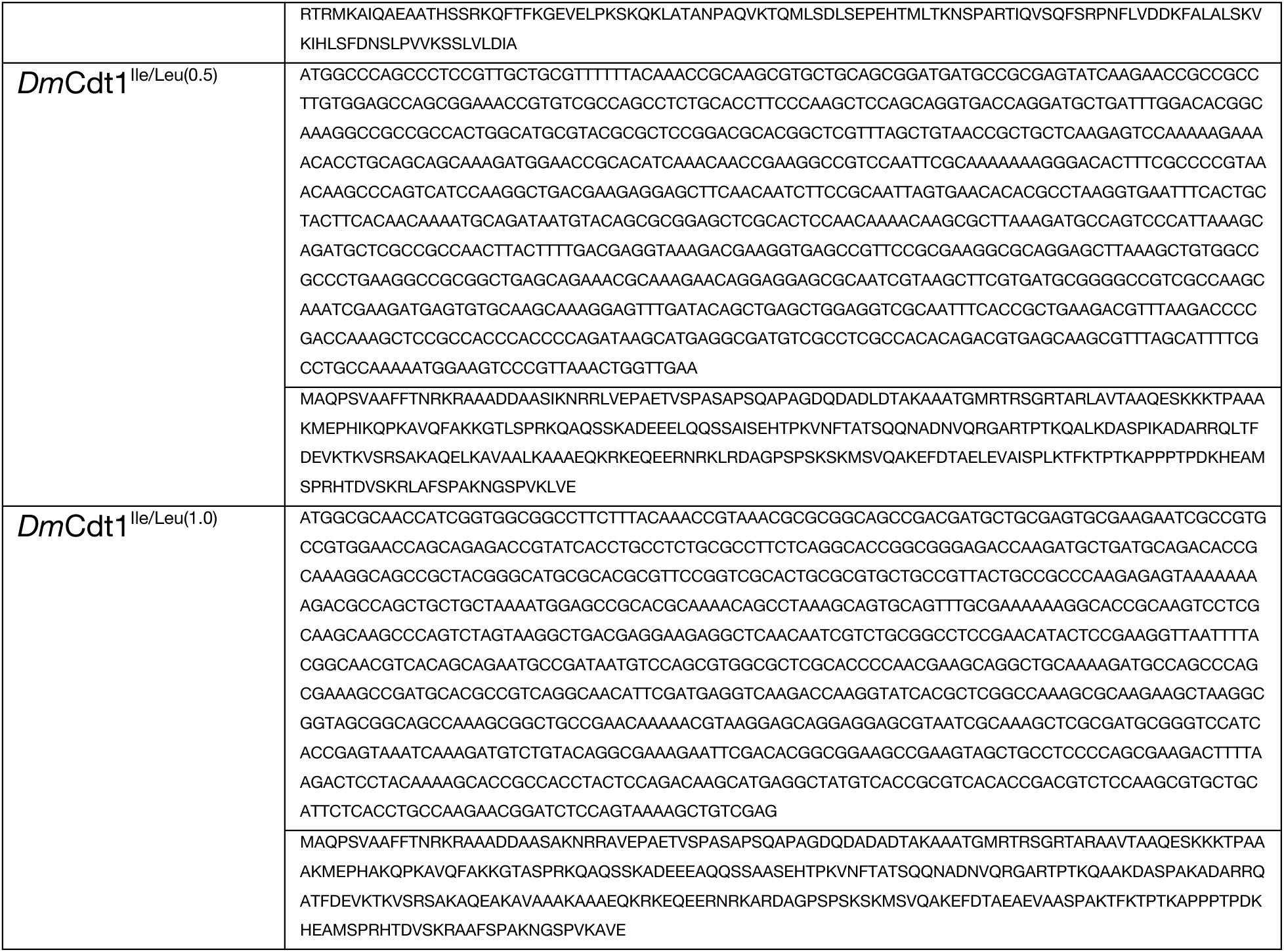
IDR sequences of the proteins used in this study.

Cell pellets were resuspended in Lysis Buffer (20 mM Tris pH 7.5, 500 mM NaCl, 30 mM Imidazole, 10% glycerol, 200 *µ*M PMSF, 1x cOmplete EDTA-free Protease Inhibitor Cocktail (Sigma-Aldrich), 1 mM BME and 0.1 mg/mL lysozyme) and lysed by sonication. Lysates were clarified by centrifugation at 30,000 xg for 1 hr and filtered through a 0.45 µm bottle-top filter unit (Nalgene Rapid-Flow, ThermoFisher). Lysates were then passed over a 5 mL HisTrap HP column (GE Healthcare) and washed with 10 column volumes (CV) of Nickel Wash Buffer (20 mM Tris pH 7.5, 1 M NaCl, 30 mM Imidazole, 10% glycerol, 200 *µ*M PMSF, 1 mM BME) followed by 5 CV of Low Salt Nickel Wash Buffer (20 mM Tris pH 7.5, 150 mM NaCl, 30 mM Imidazole, 10% glycerol, 200 *µ*M PMSF and 1 mM BME). Protein was eluted from the column with 5 CV of Nickel Elution Buffer (20 mM Tris pH 7.5, 150 mM NaCl, 500 mM Imidazole, 10% glycerol, 200 *µ*M PMSF and 1 mM BME). Eluted protein was then loaded onto a 5 mL HiTrap Heparin HP column (GE Healthcare), washed with 5 CV of Heparin Wash Buffer (20 mM Tris pH 7.5, 150 mM NaCl, 10% glycerol, 200 *µ*M PMSF and 1 mM BME) and eluted with a linear gradient of increasing salt from 150 mM – 1 M NaCl (20 mM Tris pH 7.5, 10% glycerol, 200 *µ*M PMSF and 1 mM BME). Fractions containing Cdt1 were pooled and TEV protease (QB3 Macrolab) added at a 1:20 ratio of TEV:protein and incubated overnight at 4°C. A second nickel affinity step was used to remove TEV, uncleaved protein, and the free His6-MBP tag. Finally, the sample was concentrated to 2 mL and loaded onto a HiPrep 16/60 Sephacryl S-300 HR column (GE Healthcare) equilibrated and run in Sizing Buffer (50 mM HEPES pH 7.5, 300 mM KGlutamate, 10% glycerol, 1 mM BME). Peak fractions were collected, concentrated in a 30K Amicon Ultra-15 concentrator (Millipore), flash frozen in liquid nitrogen, and stored at -80°C. The same protocol was used to purify GFP-tagged Cdt1 with the exception that TEV protease was not added and the second nickel affinity step was omitted.

*D. melanogaster* ORC and Cdc6 were purified as previously described (Parker et al., 2019).

### Fluorescence microscopy

All microscopy assays with fluorescently-labled DNA utilized “Cy5- dsDNA”, a sixty base pair duplex of sequence:

5’- GAAGCTAGACTTAGGTGTCATATTGAACCTACTATGCCGAACTAGTTACGAGCTATAAAC-3’ that had a 5’ Cy5 label (IDT or Eurofins). Duplex DNA was annealed at 100 *µ*M in 50 mM Tris pH 7.5, 50 mM KCl. Samples were prepared by mixing protein (10 *µ*M Cdt1 and Cdc6 and 5 *µ*M ORC unless otherwise noted in the text) prepared in Protein Buffer (50 mM HEPES pH 7.5, 300 mM KGlutamate, 10% glycerol, 1 mM BME) with an equal volume of equimolar Cy5-dsDNA prepared in Dilution Buffer (50 mM HEPES pH 7.5, 10% glycerol, 1 mM BME), were mixed thoroughly by pipetting and incubated for 2 mins. 7 µL of sample was spotted onto a glass slide, a coverslip placed on top, and imaged with a 60x oil objective using confocal fluorescence microscopy. Images were processed in FIJI and quantitation performed by thresholding for Cy5 signal intensity and then calculating the mean Cy5 and/or eGFP intensity within droplets. eGFP-Cdt1 phase separation was assessed in the presence of PEG-3350 (Sigma 202444-250G) by mixing equal volumes of 10 *µ*M eGFP-Cdt1 in Protein Buffer and 8% PEG dissolved in Dilution Buffer. Experiments were completed in triplicate and at least three fields of view imaged for each experiment.

### Protein depletion LLPS assay

All depletion assays with DNA utilized “dsDNA”, a sixty base pair duplex with the same sequence as Cy5-dsDNA. Duplex DNA was annealed at 100 *µ*M in 50 mM Tris pH 7.5, 50 mM KCl. Samples were prepared by mixing protein (4 *µ*M Cdt1 unless otherwise noted in the text) prepared in Protein Buffer (50 mM HEPES pH 7.5, 300 mM KGlutamate, 10% glycerol, 1 mM BME) with an equal volume of equimolar dsDNA prepared in Dilution Buffer (50 mM HEPES pH 7.5, 10% glycerol, 1 mM BME), were mixed by gently flicking the tube and incubated for 30 mins. Subsequently, the dense phase separated material was pelleted by centrifugation (10 min at 16,000 xg) and the supernatant removed to assess for protein retention by SDS-PAGE analysis and Coomassie staining. Every assay included a load control that had not been centrifuged. Samples were run on 4-20% gradient gels (BioRad 4561096).

The same protocol was used for comparing the efficiency of different polyanionic scaffolds at inducing Cdt1 phase separation except that 0.06 mg/mL of each polyanion was mixed with an equal volume of 4 *µ*M Cdt1. The polyanions included dsDNA (same sequence as above), ssDNA (same sequence as dsDNA), heparin (Sigma H3149), polyglutamate (Sigma P4886), as well as dsRNA prepared using the MEGAscript RNAi Kit (Life Technologies) from a 422 base pair PCR fragment derived from the *D. melanogaster* DPOA2 gene. For depletion assays completed in the presence of crowding reagent, PEG-3350 (Sigma 202444-250G) was used in all cases and phase separation assessed by mixing equal volumes of 16 *µ*M Cdt1 in Protein Buffer and PEG-3350 dissolved in Dilution Buffer to reach a final concentration of 0-12.5% PEG-3350. In these experiments, the samples were diluted 4-fold with water prior to analysis by SDS-PAGE to reduce migration artifacts induced by high concentrations of PEG-3350. For depletion assays with monovalent and multivalent phosphate counterions, concentrated stocks of monobasic, dibasic and tribasic potassium phosphate were prepared in Dilution Buffer and the pH adjusted to 7.5 prior to use. Experiments were completed in triplicate and band quantitation was completed using FIJI (Schindelin et al., 2012).

### DNA-binding electrophoretic mobility shift assays (EMSAs)

DNA-binding assays were completed with dsDNA containing a 5’ IRDye800 label. Protein was titrated from 16 nM to 1 *µ*M in the presence of 20 nM IRDye800-dsDNA in Assay Buffer (50 mM HEPES pH 7.5, 150 mM KGlutamate, 10% glycerol, 1 mM BME). The samples were mixed by pipetting and incubated for 45 mins at room temperature. 5 µL of each sample was run on a 7.5% PAGE gel (BioRad 4561026) pre-run at 100 volts for 1 hour and gels imaged on an Odyssey imaging system (LI-COR) through detection of IRDye800.

### Sequence analysis and bioinformatics

Amino acid heatmaps and sequence complexity were calculated in Python using the Seaborn and SciPy (Virtanen et al., 2020) modules, respectively. EMBOSS dotmatcher was used to generate sequence dotplots with parameters “Window size” = 15 and “Threshold” = 32 (Rice et al., 2000). Coiled-coil predictions were made using the Parcoil2 server (McDonnell et al., 2006).

A custom Python script was used to calculate proteome-wide statistics on ordered and disordered domains (https://github.com/mwparkerlab/Parkeretal2021_Cdt1LLPS.git). Briefly, IUPRED2 (Mészáros et al., 2018) disorder prediction was run on every protein in the *D. melanogaster* reference proteome (release UP000000803_7227.fasta) and the per-residue disorder scores were smoothed by applying a moving average over a twenty-residue window. Sequences of contiguous disorder and order longer than 100 amino acids were extracted from the data and the sequence hydropathy (Kyte and Doolittle, 1982) and amino acid composition were calculated for each region. The average fraction of each of the twenty amino acids in all ordered and disordered domains was calculated from the composition of all ordered and disordered domains longer than 100 residues. Sequence hydropathy and composition were separately calculated for the initiator IDRs, and included Orc1 residues 187-549, Cdc6 residues 1-246 and Cdt1 residues 1-297, resulting in three values that were averaged to generate initiator hydropathy and initiator composition. The difference in amino acid composition between initiators and all other ordered and disordered protein regions proteome-wide was calculated by subtracting the proteome-wide values from the initiator values.

### Analytical size exclusion and heparin affinity chromatography

To assess how charged residue distribution impacts the conformation of the Cdt1 IDR, we compared the size exclusion chromatography elution profiles of Cdt1 and Cdt1^Uniform^. 50 µL of 6 *µ*M Cdt1 or Cdt1^Uniform^ was prepared in Protein Buffer and then loaded and run on a Superose 5/150 GL sizing column (GE Healthcare) pre-equilibrated and run in Protein Buffer. We compared the heparin binding profiles of Cdt1, Cdt1^Ile/Leu(0.5)^ and Cdt1^Ile/Leu(1.0)^ by injecting 50 µL of 8 *µ*M protein in Protein Buffer onto a 1 mL HiTrap Heparin HP column (GE Healthcare) and eluting with a 10 CV linear gradient from 150 mM – 1 M NaCl in 20 mM Tris pH 7.5, 10% glycerol, 1 mM BME.

All IDR sequences were fused back to wild-type Cdt1 residues 298-743.

## ACKNOWLEDGEMENTS

We thank past and present members of the Berger and Botchan labs for helpful discussion and advice. This work was supported by an NIH NRSA postdoctoral fellowship (F32GM116393, to MWP), by the UC Berkeley Jesse Rabinowitz Award (to JAK) and by the NCI (R01CA030490, to JMB and MRB).

## AUTHOR CONTRIBUTIONS

Conceptualization, M.W.P., M.R.B. and J.M.B.; Methodology, M.W.P.; Investigation, M.W.P., A.H. and J.A.K.; Writing - Original Draft, M.W.P, M.R. and J.M.B.; Writing - Review & Editing, M.W.P., A.H., J.A.K., M.R.B. and J.M.B.; Visualization, M.W.P.; Funding Acquisition, M.W.P., M.R.B., J.M.B.; Supervision, M.W.P., M.R.B. and J.M.B.

## SUPPLEMENTAL INFORMATION TITLES AND LEGENDS

**Figure 1 – figure supplement 1:** *D. melanogaster* ORC and Cdc6 phase separation is resistant to treatment with 1,6-hexanediol. A) 2.5 *µ*M ORC was combined with 2.5 *µ*M Cy5-dsDNA and phase separation assessed by confocal fluorescence microscopy in the absence (left) and presence (right) of 10% 1,6-hexanediol. B) 5 *µ*M Cdc6 was combined with 5 *µ*M Cy5-dsDNA and phase separation assessed by confocal fluorescence microscopy in the absence (left) and presence (right) of 10% 1,6-hexanediol.

**Figure 3 – figure supplement 1:** Dotplot analysis of various phase-separating IDRs to identify the presence of repetitive motifs. EMBOSS Dotmatcher (Madeira et al., 2019) was used to assess the presence of repetitive elements in the IDRs of A) Laf1, B) Ddx4, C) hnRNPA1, D) RPB1, E) MED1, and F) BuGZ. All proteins analyzed contain short repetitive motifs.

**Figure 3 – figure supplement 2:** Cdt1^ScrIDR^ undergoes phase separation in the presence of DNA and can recruit wild-type Cdt1. A) 5 *µ*M Cdt1^ScrIDR^ was combined with 5 *µ*M Cy5-dsDNA and phase separation assessed by fluorescence microscopy. B) Droplets were prepared with 5 *µ*M Cy5-dsDNA and either 5 *µ*M wild-type Cdt1 (left two columns) or 5 *µ*M Cdt1^ScrIDR^ (right two columns) and 1 *µ*M eGFP-Cdt1 was subsequently added to each reaction. Samples were imaged by fluorescence microscopy. C) Quantitation of panel B). The level of eGFP-Cdt1 recruitment was assessed by calculating green signal intensity within the droplets containing Cy5-dsDNA and either Cdt1 or Cdt1^ScrIDR^.

**Figure 5 – figure supplement 1:** Heparin binding propensity of Cdt1 and the Cdt1 branched hydrophobic residue variants, Cdt1^Ile/Leu(0.5)^ and Cdt1^Ile/Leu(1.0)^. Cdt1 (top) and Cdt1 variants (middle and bottom) were applied to a heparin column and eluted off with a linear gradient of 150 mM – 1 M NaCl. All three proteins elute off at similar salt concentrations.

**Figure 5 – figure supplement 2:** Heatmap analysis of the amino acid composition of initiator IDRs (Cdt1, Orc1, Cdc6) and of an internal IDR from Chiffon, the fly homolog of Dbf4. These four proteins have similar amino acid sequence signatures, suggesting that Chiffon may undergo a similar DNA-dependent phase separation reaction or may co-partition into initiator condensates.

