## Supplementary figures and images for "Molecular determinants of phase separation for *Drosophila* DNA replication licensing factors"

### Supplemental Figures

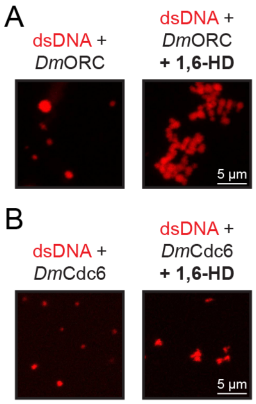

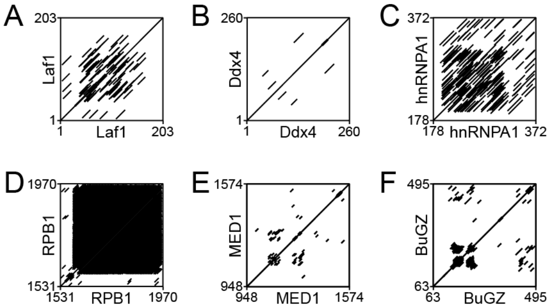

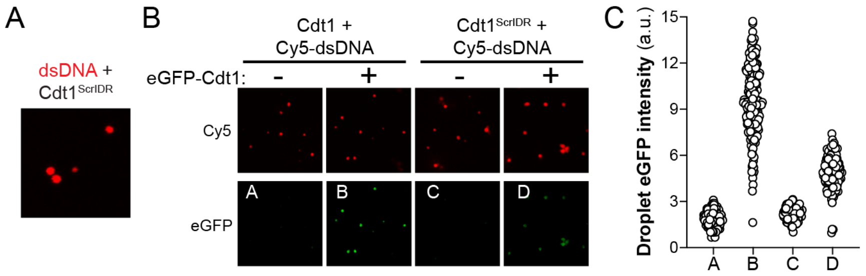

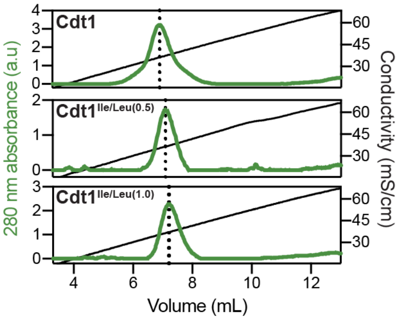

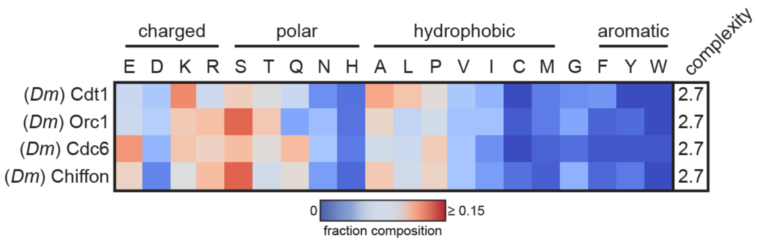
